# Temporal dependence of shifts in mu opioid receptor mobility at the cell surface after agonist binding observed by single-particle tracking

**DOI:** 10.1101/402826

**Authors:** Marissa J. Metz, Reagan L. Pennock, Diego Krapf, Shane T. Hentges

## Abstract

Agonist binding to the mu opioid receptor (MOR) results in conformational changes that allow recruitment of G-proteins, activation of downstream effectors and eventual desensitization and internalization, all of which could affect receptor mobility. The present study employed single particle tracking (SPT) of quantum dot labeled FLAG-tagged MORs to examine shifts in MOR mobility after agonist binding. FLAG-MORs on the plasma membrane were in both mobile and immobile states under basal conditions. Activation of FLAG-MORs with DAMGO caused an acute increase in the fraction of mobile MORs, and free portions of mobile tracks were partially dependent on interactions with G-proteins. In contrast, 10-minute exposure to DAMGO or morphine increased the fraction of immobile FLAG-MORs. While the decrease in mobility with prolonged DAMGO exposure corresponded to an increase in colocalization with clathrin, the increase in colocalization was present in both mobile and immobile FLAG-MORs. Thus, no single mobility state of the receptor accounted for colocalization with clathrin. These findings demonstrate that SPT can be used to track agonist-dependent changes in MOR mobility over time, but that the mobility states observed likely arise from a diverse set of interactions and will be most informative when examined in concert with particular downstream effectors.

## Introduction

The mu opioid receptor (MOR) is a G-protein coupled receptor (GPCR) responsible for many of the physiological effects of endogenous opioids, as well as clinically important exogenously administered opioids (e.g. morphine, codeine, fentanyl). Most functional studies of the MOR have focused on the output produced by the inhibition or activation of effectors such as adenylyl cyclase or G-protein coupled inwardly rectifying K^+^ channels, as this generally provides a good proxy for signaling through the receptor and for receptor desensitization. However, effector-based assays often report the mean response across the population of receptors and can have limited spatial and temporal resolution. Further, readouts for each effector of interest can have distinct sensitivity and resolution, and assays for each effector are usually performed separately from one another. Single-molecule imaging and tracking of MORs may provide a direct measure of receptor activation and desensitization that can accompany population-based assays and may help uncover underlying heterogeneity in the overall population of receptors that is difficult to detect with traditional assays.

Imaging-based studies of directly labeled MORs have provided an alternative to effector-dependent assays and have been used to examine MOR mobility^1-7^ and subcellular localization^3,8-10^. Studies of MOR mobility utilizing fluorescence recovery after photobleaching (FRAP) and fluorescence correlation spectroscopy (FCS) have shown that MORs can exist in a variety of mobility states on the membrane. However, existing studies suggest that agonist binding either increases^4,7^ or decreases^5^ lateral mobility of MORs. These conflicting results may be due to the use of different cell lines, different experimental conditions (e.g. temperature and agonist exposure duration) and the fact that these experiments relied on bulk approaches that reflect average mobility of the receptors as a population.

Single-particle tracking (SPT) approaches allow for examination of individual receptor behavior without reliance on effector-based readouts. Further, single-molecule techniques allow the evaluation of heterogeneities and molecule-to-molecule variations without relying on *a priori* models as required in ensemble measurements, such as FRAP^11,12^. Specific mobility states of individual receptors detected with SPT may correlate with distinct interactions with select effectors and other binding partners. Prior studies examining MOR mobility using SPT have shown differing results as to what these distinct mobility states are. One study tracking the MOR reported that most receptors are confined within mobile microdomains, while a smaller fraction of receptors exhibit slow, directed diffusion^2^. Another study reported short-term confinement within specific membrane compartments followed by diffusion of the receptor between these compartments^6^. The relationship between mobility states, agonist binding, and receptor-effector interactions have yet to be examined using such assays.

The present study was designed to determine if SPT could provide a reliable approach to detect agonist-induced changes in mobility of the MOR that correspond to interactions with G-proteins and recruitment to clathrin coated pits (CCPs) subsequent to agonist binding. Tracking of the MOR was performed in AtT20 cells stably expressing a FLAG-epitope-tagged MOR construct (FLAG-MOR)^13^ conjugated with quantum dots (Qdots) via an anti-FLAG antibody. Signaling through MORs has been extensively characterized in AtT20 cells^13-15^. MORs in AtT20 cells couple to endogenously-expressed GIRKs^14,16^, P/Q-type VDCCs^13^, adenylyl cyclase^17^, and G-protein coupled receptor kinases^14,18^ via pertussis-toxin sensitive Gαi/o proteins^19^. Further, AtT20 cells stably expressing FLAG-MORs exhibit relatively consistent and moderate expression of the receptor and display activation, desensitization, and internalization phases similar to MORs in neurons^13^. Thus, AtT20 cells stably transfected with Flag-MOR provide a good system for examining real-time dynamics of the MOR in the plasma membrane before and during agonist exposure.

Here, MOR mobility was investigated in response to two different drugs: morphine, a low efficacy agonist that induces very little internalization, and DAMGO, a high efficacy agonist known to induce internalization. Mobility in response to these drugs was investigated after 1 minute or 10 minutes of agonist exposure. These time points were selected because 1 minute of agonist exposure has been shown to induce receptor coupling to G-proteins and signaling through various effectors, whereas 10 minutes of agonist exposure can lead to receptor desensitization and internalization in an agonist-dependent manner^13^. Quantitative diffusion analysis of trajectories of Qdot 565-conjugated FLAG-MORs shows that receptors can be found in both immobile and mobile states under basal conditions. Activation of FLAG-MORs with DAMGO, but not morphine, caused an initial increase in the population of mobile receptors (at 1 minute of exposure), followed at later time points by a decrease in the mobile fraction (10 min agonist exposure). These findings show that activation of the FLAG-MOR does not result in a uniform decrease in the mobility of the receptors, but instead it dynamically changes the fraction of FLAG-MORs found in either more mobile or immobile states depending upon time observed after agonist exposure and the agonist applied. Further, investigation of receptors in the mobile state revealed transient confinement partially influenced by G-protein binding. However, inspection of tracks after 10 min of DAMGO exposure revealed that the mobility state was a poor proxy for colocalization with clathrin. The results indicate that ensemble measurements of receptor mobility, such as average mean square displacement (MSD), may not capture the full extent of receptor activity, and that single-particle tracking can better account for heterogeneity in signaling states. Altogether, mobility state can provide information about signaling state, but may not account for the range of potential effectors a receptor is coupled to.

## Results

In order to visualize MORs, FLAG-MORs in AtT20 cells were labeled with Qdots. Cells were imaged under differential interference contrast (DIC) to assess cell morphology (Fig. 1A) and under fluorescence using spinning disk confocal to detect Qdots (Fig. 1B). Trajectories of labeled FLAG-MOR were obtained as shown in the example in Fig. 1C. FLAG-MOR mobility was investigated before agonist application and after 1 or 10 minutes of treatment with agonist (10 µM). The time-averaged 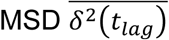 of individual Qdot-MOR trajectories were computed for time lags *t_lag_* up to 2.5 s for each experimental condition (Eq. 1 in methods section). Fig. S1 shows a representative set of individual MSDs for trajectories in various conditions. Most trajectories exhibit sub-diffusive behavior, i.e., the MSD of individual trajectories is not linear in lag time but it scales as a power law 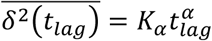, with α < 1 where α.is the anomalous exponent and *K*_*α*_ is the generalized diffusion coefficient20-22.Furthermore, a substantial number of trajectories have a flat MSD, indicating that these particles are either immobile or confined to small domains^23^. In all cases, only particles that were observed for at least 3 s were analyzed. Clear differences were observed between the MSD under basal conditions and those in cells treated with DAMGO or morphine (Fig. 2A). Short DAMGO treatment (1 min) increased the anomalous exponent *α* of the average MSD from 0.49 ± 0.12 in the no-drug condition to 0.55 ±0.02. No change was observed from no drug in the anomalous exponent of the average MSD after 1 min morphine (0.48 ± 0.04, *p* = 0.88). In contrast, longer DAMGO treatment (10 min) decreased the anomalous exponent of the average MSD to 0.38 ± 0.02 (*p* = 0.014 compared to no-drug), as did 10 min morphine treatment (0.40 ± 0.02, *p* = 0.02). Further, the generalized diffusion coefficient, *K*_α_, of the averaged MSD increased after the 1 min DAMGO exposure from 0.087 ± 0.005 μm^2^/s^0.49^ (no-drug condition) to 0.104 ± 0.006 μm^2^/s^0.55^ (*p* = 0.034). *K*_α_ also increased after 1 min morphine treatment (0.16 ± 0.03 μm^2^/s^0.48^, *p* = 0.002) as well as after 10 min morphine treatment (0.14 ± 0.02 μm^2^/s^0.40^, *p* = 0.003). Longer DAMGO exposure (10 min) decreased the generalized diffusion coefficient to 0.057 ± 0.004 μm^2^/s^0.38^ (*p* = 0.0004 compared to no-drug). Together, these results indicate that acute agonist exposure increases the average diffusivity of the receptor and this average diffusivity can either increase or decrease after prolonged agonist exposure.

**Figure 1.**
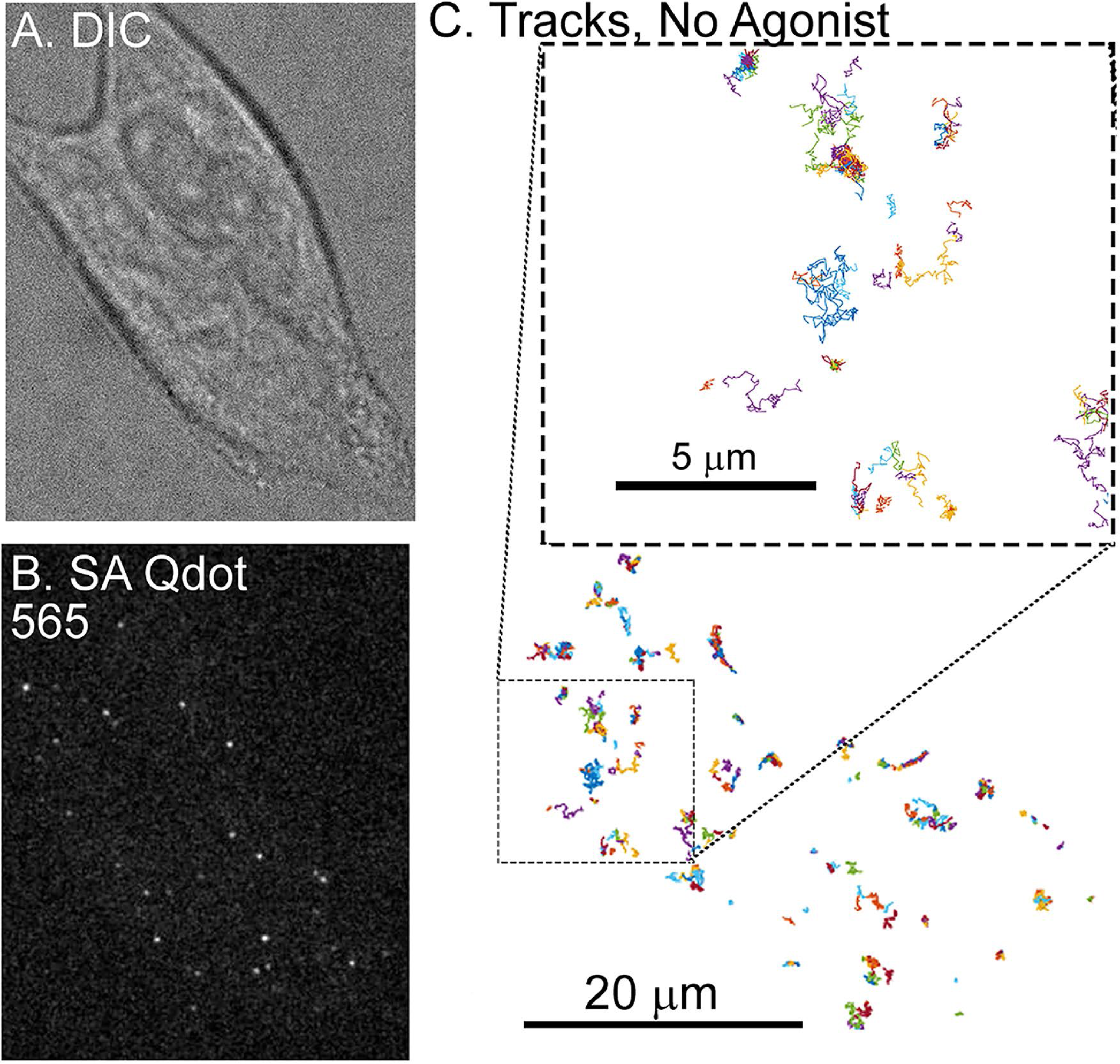
Single-particle tracking of FLAG-MORs in AtT20 cells. **A)** Representative DIC image of an AtT20 cell and (**B)** fluorescent image of SA Qdot 565 conjugated FLAG-MORs in the same cell. Labeling density in this image is typical for experiments carried out in this study. **C)** Trajectories of SA Qdot 565-conjugated FLAG-MORs imaged in the absence of drug from the cell shown in (A) and (B). All of the displayed trajectories were obtained from a single 1000-frame video segment. The upper square in (C) shows an enlarged image of trajectories from the area indicated by the lower dashed box. Both mobile and immobile trajectories can be seen in the inset, as well as confined and free portions of mobile trajectories.

**Figure 2.**
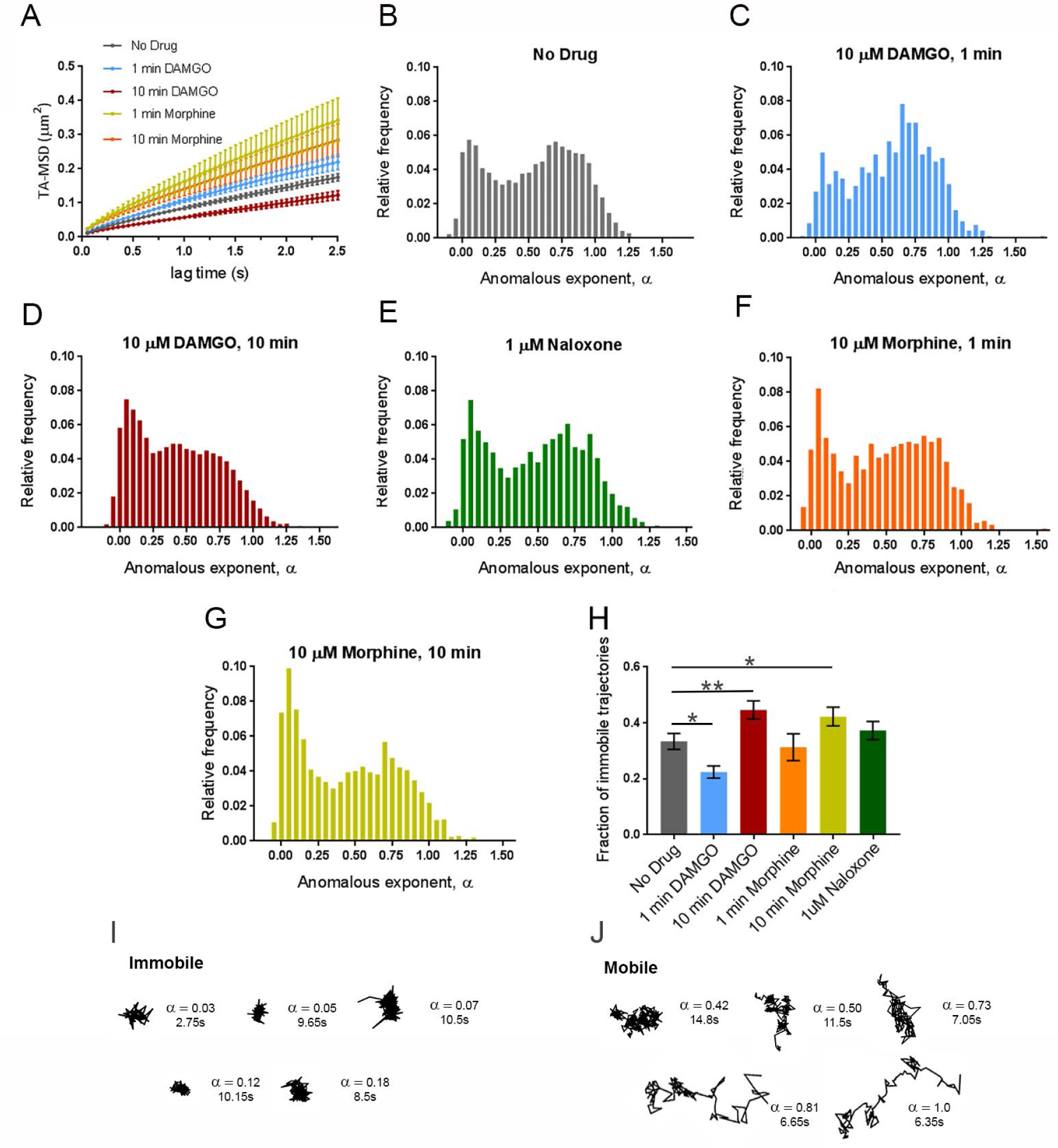
Dynamic changes in MOR mobility exist depending upon the time of drug application. **A)** Time averaged MSD averaged over all tracks for each experimental condition showing increases in the MSD with 1 min DAMGO, 1 min morphine, and 10 min morphine treatments and a decrease with 10 min DAMGO treatment. Distribution of anomalous exponents α, obtained from the TA-MSD of individual trajectories for tracks in the **B)** no drug (n=17 cells, 8465 trajectories), **C)** 1 min DAMGO (n=12 cells, 930 trajectories), **D)** 10 min DAMGO (n=13 cells, 8465 trajectories), **F)** 1 min morphine (n=8 cells, 882 trajectories), **G)** 10 min morphine (n=10 cells, 2415 trajectories), **E)** and Naloxone (n=7 cells, 8227 trajectories) condition. All conditions exhibit bimodal distributions with peaks centered at *α* = 0.09 and *α* = 0.71 for immobile and mobile populations, respectively. **H)** Compared to the no drug condition, the population of *α* < 0.27 decreases in the 1 min DAMGO condition, increases in the 10 min DAMGO and 10 min morphine conditions, and does not change after 1 min morphine or Naloxone application. Example trajectories from **I)** immobile (α < 0.27) and **J)** mobile (α > 0.27) populations are shown with their corresponding α values and lengths. \**p <* .05, ** *p* < .01, Kruskall-Wallis one-way ANOVA with multiple comparisons using Bonferroni correction, error bars represent SEM for cells.

Ensemble measurements, such as average MSD, however, may mask important characteristics in the data that can be revealed by measuring individual trajectories. Thus, the anomalous exponent *α* was computed for individual trajectories in each experimental condition. Very low anomalous exponents are indicative of particles that explore a more compact region and present a nearly flat MSD while larger exponents indicate that the trajectory explores a less compact region and present an MSD that increases with lag time^24^. The histogram for the no-drug condition shows a large scattering of *α* values, indicating marked heterogeneity in the diffusion behavior of FLAG-MORs (Fig. 2B). In particular, two populations are clearly visible, one with a narrow distribution centered at *α* = 0.09 and another with a broader distribution centered at *α* = 0.71. These populations can be separated by using a threshold in *α*. We selected a threshold *α =* 0.27, based on the local minimum that is found in the histogram between the two populations. Examination of the tracks within these two populations shows relatively more confinement in the low *α* population, and relatively less confinement in the high *α* population (see Fig. 2I & J for example tracks). Henceforth, tracks with *α* < 0.27 are referred to as immobile and tracks with *α* ≥.0.27 are referred to as mobile. The distribution of *α* values is significantly altered by 1-min DAMGO (Fig. 2C), 10-min DAMGO (Fig. 2D), and 10-min morphine (Fig. 2G) exposure compared to the no-drug condition (Fig. 2B). The fraction of immobile tracks in each condition is shown in Fig. 2H. In the no drug condition, the immobile fraction is 0.33 ± 0.03. This decreased to 0.23 ± 0.02 in the 1-min DAMGO condition (*p* = 0.016) and increased after 10 min of DAMGO exposure to 0.45 ± 0.03 (*p* = 0.009). Morphine exposure for 1 min does not cause a statistically significant decrease in the immobile fraction (0.31 ± 0.05; *p* = 0.842 compared to no-drug, see Fig. 2F for distribution), but like 10 min DAMGO, 10 min of morphine exposure exhibits a higher immobile fraction at 0.42 ± 0.03 (*p* = 0.047). The difference between 1 min DAMGO and 1 min morphine might be due to the lower efficacy of morphine, differing effector coupling, or a differing time course of activation induced by the two agonists. Regardless, it is clear that acute exposure to DAMGO reduces the fraction of immobile receptors, suggesting that mobility increases when the receptor is actively signaling. As expected, there is no significant change in the fraction of immobile trajectories caused by the neutral antagonist Naloxone (0.37 ± 0.03, *p* = 0.147 compared to the no drug condition, see Fig. 2E for distribution), indicating that ligand binding alone was not sufficient to cause the altered mobility observed.

Despite the advantages of Qdot labeling in terms of brightness and photostability^25-27^, quantum dots have the potential to reduce surface protein mobility^28^. Further, antibody-based detection has the potential to cause receptor crosslinking. To address these concerns, we compared our results with Qdot-labeled MORs to measurements of Flag-MOR bound to the fluorescent agonist Dermorphin-488^29^. Exposure to Dermorphin-488 (10 min) resulted in *α* values similar to those obtained after 10 min of DAMGO exposure in cells with Qdot-labled MORs (Fig. S2C). Further, if Qdots were crosslinked during tracking, the intensity of a measured particle would correspond to slower diffusion, as aggregates of Qdots would be expected to be both brighter and slower. This is not the case, however, as the MSD is not correlated with the intensity of each particle (Fig. S2D). Thus, we exclude Qdot artifacts and crosslinking as the root for the immobilization observed. Overall, the findings indicate that the MOR shows dynamic alterations in mobility depending upon whether it is observed 1 or 10 min after drug treatment.

Since there was a shift towards increased mobile receptors (reduced fraction of immobile tracks) with 1 min DAMGO exposure (Fig. 2H) when signaling is expected to be high, we next focused on the mobile tracks specifically and limited the analysis to DAMGO since morphine did not result in a significant shift in the mobile fraction at 1 min. When comparing the MSDs of mobile trajectories (Fig. 3A), the anomalous exponent of the average MSD in the no drug condition was 0.62 ± 0.01 and was similar (0.65 ± 0.02) after 1 min of DAMGO. However, 10 min of DAMGO exposure decreased the anomalous exponent of the average MSD to 0.54 ± 0.03 (*p* = 0.02). Further, the *K*_α_ of the averaged MSD for mobile trajectories was 0.11 ± 0.02 μm^2^/s^0.62^ in the no drug condition and 0.12 ± 0.03 μm^2^/s^0.65^ in the 1 min DAMGO condition,whereas it was 0.087 ± 0.019 μm^2^/s^0.54^ in the 10 min DAMGO condition (p = 0.041, no drug compared to 10 min DAMGO). This result suggests that, even within the mobile population of receptors, prolonged agonist exposure may lead to relatively reduced mobility.

**Figure 3.**
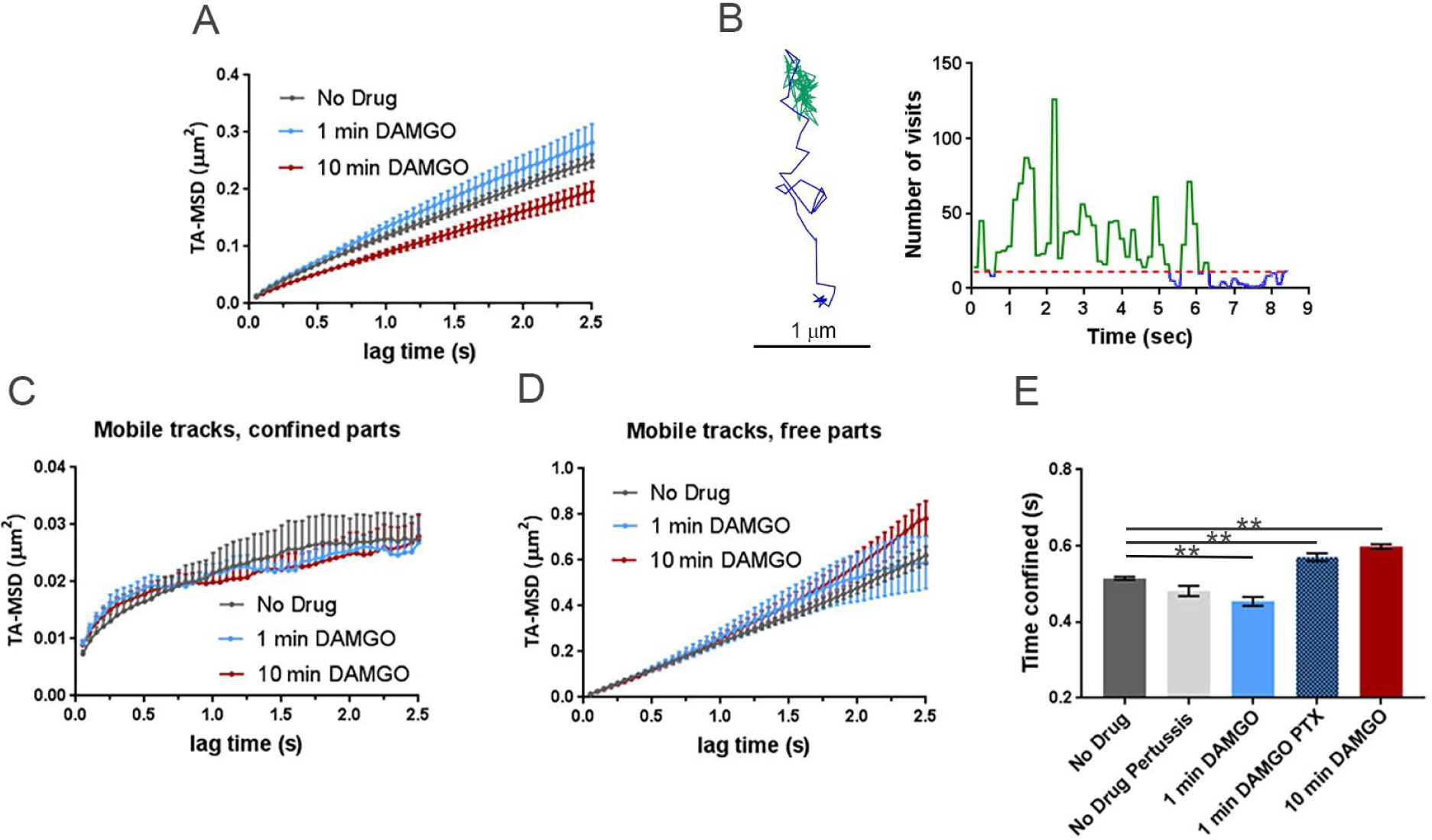
Differences between MSD of mobile tracks can be explained by dwell times between experimental conditions and G-protein binding. **A)** MSDs of mobile trajectories decreased after 10 min DAMGO treatment (*p* = 0.031). No drug (n=5855 trajectories), 1 min DAMGO (n= 697 trajectories), 10 min DAMGO (n=3958 trajectories). **B)** Example of a mobile track separated into free (blue) and confined (green) parts based on recurrence analysis (see materials and methods). A threshold value of 11 visits inside the circle was used to define confined portions (red dotted line). The right panel shows a time series of the number of visits inside the circle for the example track shown, and free and confined portions are colored as in the example track. When the trajectory is segmented into **C)** confined and **D)** free portions, the MSDs are no longer different between experimental conditions (confined *p* = 0.150, free *p* > 0.99). **E)** Confined dwell times decrease after acute DAMGO treatment and increase with longer DAMGO treatment. Application of 100 ng PTX increases confined dwell times. \*\**p* < 0.01, Kruskall-Wallis one-way ANOVA with multiple comparisons using Bonferroni correction, error bars represent SEM for individual tracks.

To examine the root of this reduced mobility, we identified within the mobile trajectories (*α* > 0.27) periods of time where the molecule is in one of the two modes of motion, namely, either confined or free. To identify the times at which MOR motion changes from confined to free and vice versa, we used a recently published method based on recurrence statistics^30,31^ This methodology allows the examination of the same receptor in two different mobility states. Briefly, for each two consecutive locations in a trajectory, a circle is drawn with its center halfway between them and the number of visits within this circle is counted. Periods that undergo confinement revisit the same region many times and thus the recurrence statistic is very high^30^. However, during free periods, particle exploration is less compact and thus the recurrence statistic is low (also see materials and methods, Fig. 3B). After segmenting the mobile trajectories into free and confined states, the MSDs of free and confined states did not differ between experimental conditions (confined *p* = 0.150, free *p* > 0.99, Figs. 3C & D). To better understand this narrowing in the MSD values between experimental conditions after recurrence analysis segmentation, we measure the dwell times of trajectories within free and confined states. One minute of DAMGO treatment decreased the time mobile tracks spend confined from 0.515 ± 0.004 s to 0.454 ± 0.011 s (*p* < 0.0001, Fig. 3E, light blue bar) and 10 min DAMGO increased confined periods to 0.598 ± 0.006 s (*p* < 0.0001, Fig 3E, red bar).

The conformational shift in MORs introduced by agonist binding rapidly results in interactions with G-proteins to induce signaling. Thus, we next investigated how disrupting G-protein interactions affects the time mobile MORs spend in free and confined states to understand if confinement of mobile trajectories is associated with G-protein interaction. Cells were treated overnight with pertussis toxin (PTX, 100 ng) to inhibit G-protein/receptor interactions and dwell times in free and confined portions of mobile tracks were determined. Treatment with PTX without subsequent application of drug did not change confined dwell times of mobile trajectories (0.482 ± 0.014 s, *p* = 0.11, n = 9 cells; Fig. 3E, light gray bar). 1 min DAMGO after PTX exposure increased confined time for mobile tracks (0.570 ± 0.011 s, *p* < 0.0001) compared to no drug, n = 11 cells, Fig. 3E, darker blue bar). Interestingly, the distribution of alpha values for DAMGO (1 min)-PTX treated cells shows no difference in the immobile fraction compared to the no drug condition (0.27 ± 0.03, *p* = 0.19, data not shown).Therefore, G-protein binding appears to contribute specifically to the time mobile trajectories spend in the free state, and this is revealed using confinement analysis.

Previous work indicated that after 10 min of DAMGO exposure, half of Flag-MORs in AtT20 cells have been internalized and internalization is over half maximal^13^. Because of the observed shift towards lower *α* values after 10 min of DAMGO treatment in our study, we hypothesized that capture into CCPs accounted for this shift towards receptor immobility. Thus we examined colocalization of the MOR with GFP-labeled clathrin light chain (GFP-CLC) as previously described^27,32-34^. Representative images of GFP-CLC and MOR co-tracking can be seen in Fig. 4A along with representative tracks around GFP-CLC puncta in Figs. 4B&C. Ten minutes of DAMGO treatment increased colocalization with GFP-CLC (from 4.8 ± 0.6% to 12.1± 1.2%, *p* < 0.0001, Fig. 5A), and the average distance to GFP-CLC was reduced by drug treatment (from 1.88 ± 0.05 μm to 1.34 ± 0.04 μm, *p* < 0.0001, Fig. 5B & C). Furthermore, the distribution of distances to GFP-CLC exhibited a leftward shift in the individual track distances after 10 min DAMGO and a peak relative frequency at 0.13 and 0.75 μm for the no drug and 10 min DAMGO conditions, respectively. However, the immobile population of receptors was no more colocalized with GFP-CLC than the mobile population of receptors in the 10 min DAMGO condition (immobile 12.1 ± 1.9% vs mobile 12.1 ± 1.5%, *p* = 0.54, Fig. 6A). Distance to GFP-CLC was also unchanged between mobile and immobile populations after 10 min of DAMGO treatment (immobile 1.34 ± 0.07 μm vs. mobile 1.33 ± 0.05 μm, *p* = 0.18, Fig. 6C & D), and the distribution of distances to GFP-CLC was similar (red and green lines, Fig. 6C). Inspection of GFP-CLC colocalization within the confined and free portions of mobile receptors after 10 min of DAMGO treatment showed that neither of these populations fully accounted for GFP-CLC colocalization (confined 14.6 ± 2.6% vs. free 10.8 ± 2.0%, *p* = 0.99, Fig. 6B). Distance to GFP-CLC was also unchanged between free and confined portions of mobile tracks (confined 1.29 ± 0.07 μm vs. free 1.38 ± 0.07 μm, *p* = 0.32, Fig. 6E & F), and the distribution of distances to GFP-CLC was similar with a slight leftward shift from 1.3 (green line) to 0.73 (red line) in the 10 min DAMGO free and confined portions, respectively (Fig. 6E). Thus, mobility itself is not a good proxy for association with CCPs.

**Figure 4.**
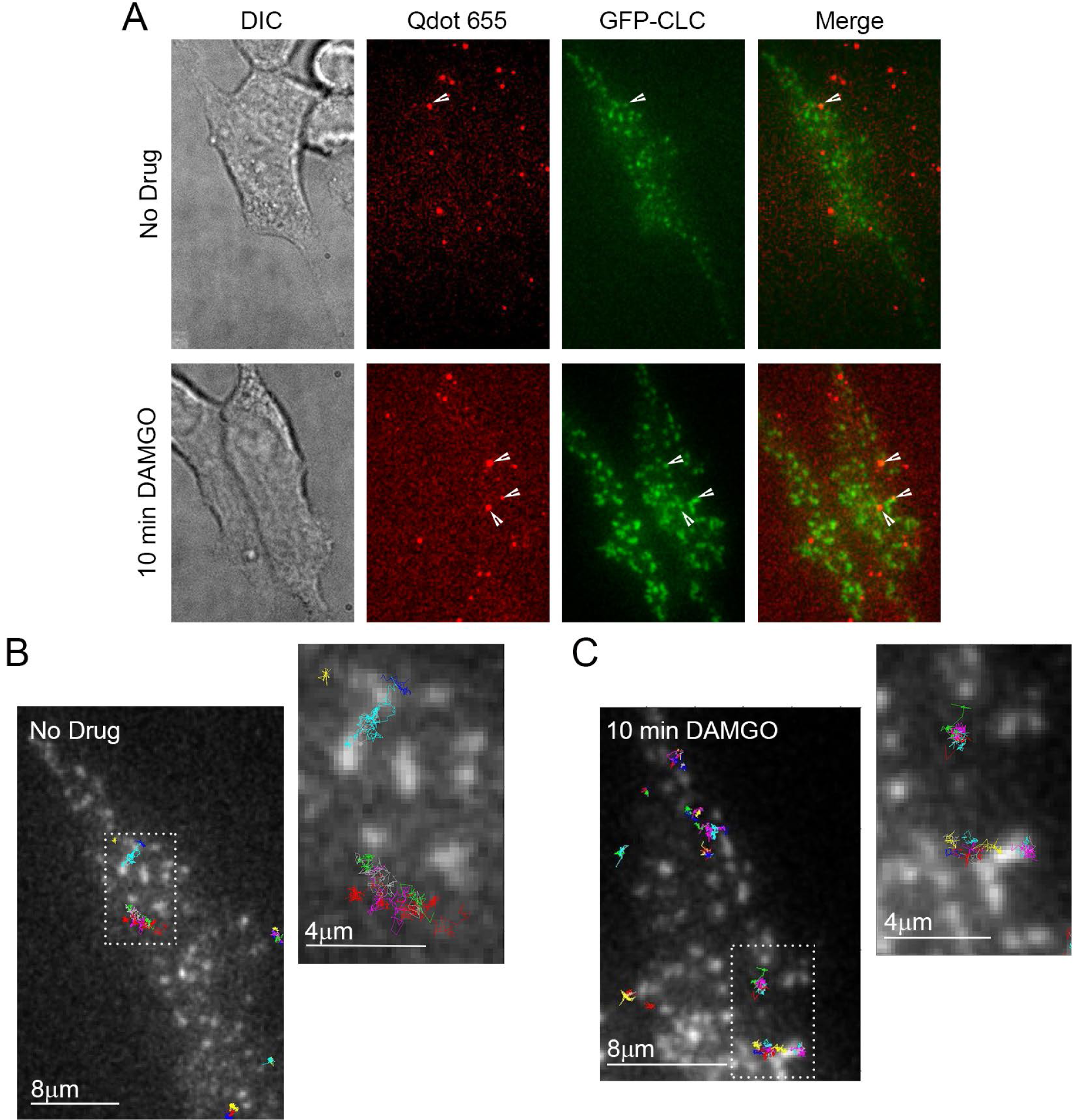
Representative GFP-CLC and Qdot655 co-tracking images. **A)** DIC and fluorescence images of cells labeled with Qdot 655-FLAG-MOR and GFP-CLC in the no drug and 10 min DAMGO conditions. White arrows indicated co-localization of Qdot-labeled FLAG-MOR and GFP-CLC. **B)** Overlay of FLAG-MOR trajectories on GFP-CLC image from an untreated cell. White dotted boxes indicate the area represented in the zoomed image. **C)** FLAG-MOR trajectories overlaid on GFP-CLC from a cell treated for 10 min with 10 αm DAMGO.

**Figure 5.**
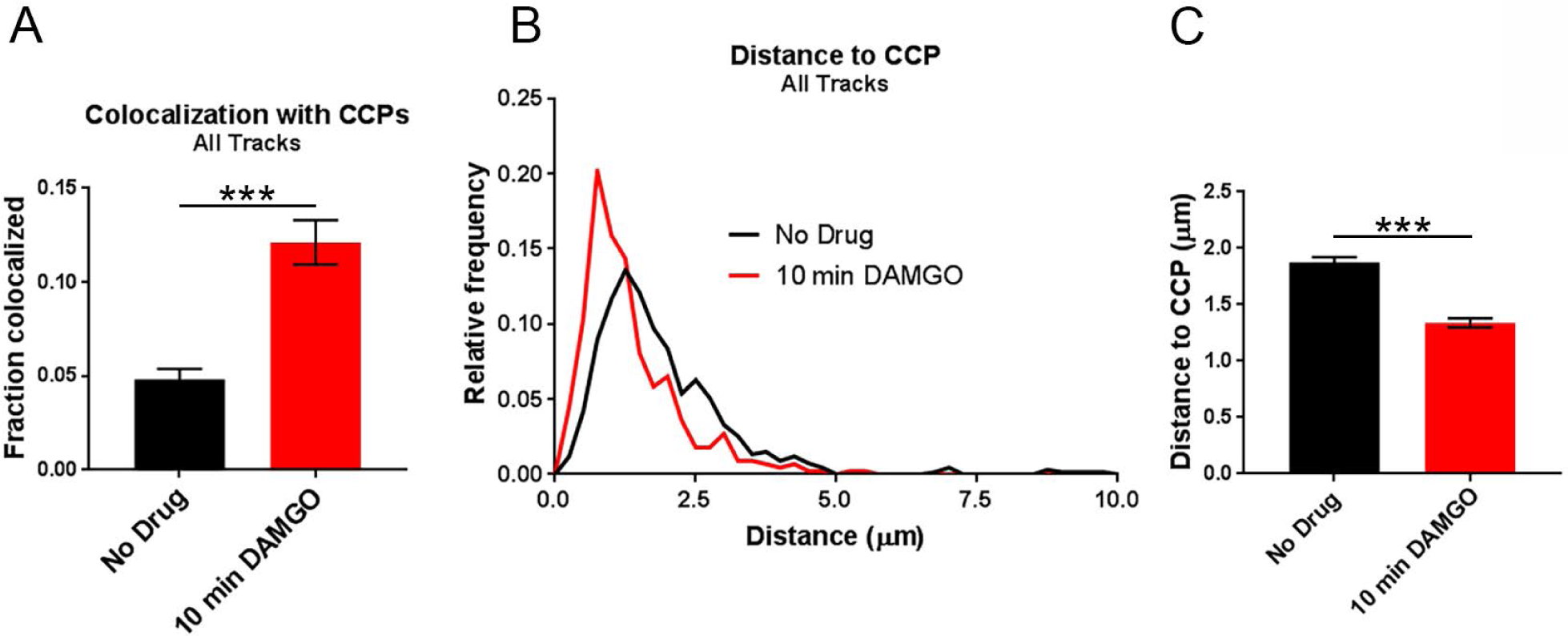
Colocalization with CCPs increases across all tracks after DAMGO (10 min) application. **A)** When comparing all tracks, colocalization with CCPs increases after 10 min of DAMGO treatment. No drug n = 9 cells, 670 trajectories; 10 min DAMGO n = 8 cells, 446 trajectories. **B)** Distribution of distances to CCPs is shifted leftward after 10 min of DAMGO treatment **C)** Combined data from (**B)** showing an overall decrease in the distance to CCPs for MORs treated with DAMGO for 10 min. ***p < 0.0001, Kruskall-Wallis one-way ANOVA with multiple comparisons using Bonferroni correction, error bars represent SEM for individual tracks.

**Figure 6.**
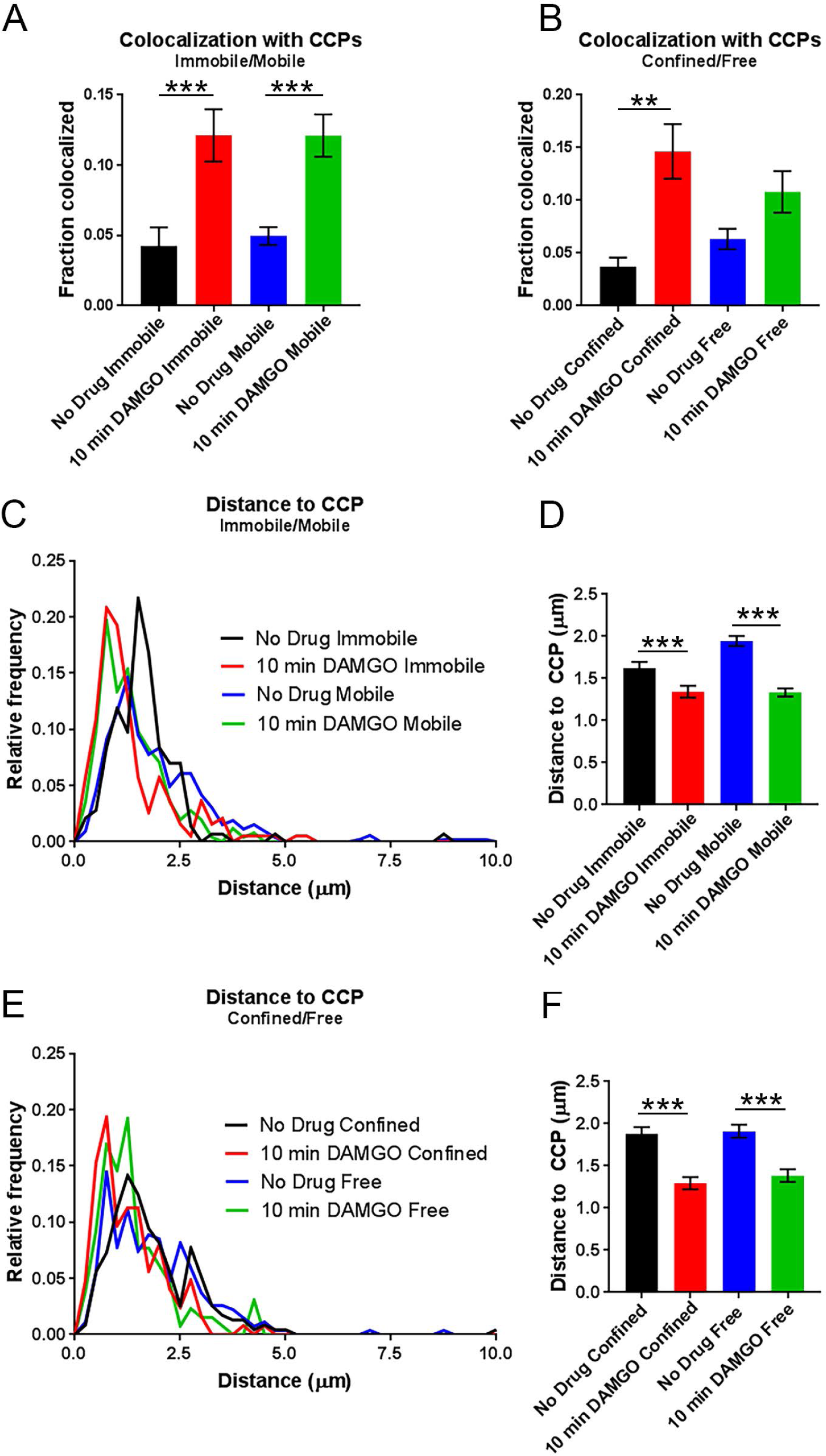
Mobility state does not reflect colocalization with clathrin. **A)** Colocalization with CCPs does not differ between immobile and mobile populations within either the no drug nor the 10 min DAMGO experimental conditions. However, 10 min DAMGO still increases the fraction of MORs colocalized with CCPs both in mobile and immobile populations compared to no drug. **B)** Colocalization with CCPs also does not differ between confined and free portions of mobile tracks within the 10 min DAMGO or no drug experimental conditions. **C)** Distance to CCPs is decreased between no drug and 10 min DAMGO experimental conditions, but is not different between immobile and mobile fractions within experimental conditions. **D)** Combined data from (**C)** shows a clear lack of difference for distance to CCPs between immobile and mobile fractions from no drug and DAMGO treated cells. **E)** Distance to CCPs is not changed between confined and free portions of mobile tracks within either no drug or 10 min DAMGO conditions.**F)** Combined data from (E). \*\**p* < 0.01, \*\*\**p* < 0.0001, Kruskall-Wallis one-way ANOVA with multiple comparisons using Bonferroni correction, error bars represent SEM for individual tracks.

## Discussion

The present single-molecule study was undertaken to characterize time-dependent changes in FLAG-MOR diffusion induced by acute and extended applications of the MOR agonists morphine and DAMGO and to understand if specific mobility states of the MOR could clearly reflect interactions with specific effectors. Under basal conditions, both mobile and immobile populations of FLAG-MORs were detected, evident in the bimodal distribution of the anomalous exponent *α* and consistent with recent work showing that MORs exist both in lipid-enriched nanodomains and more freely-distributed in the plasma membrane prior to agonist binding^35^. The bimodal distribution of FLAG-MORs observed in the present study was skewed towards more mobile tracks when cells were exposed to a maximal concentration of DAMGO (10 µM) for one minute. However, DAMGO treatment for ten minutes resulted in a dramatic increase in the proportion of immobile FLAG-MORs. Further, overall track mobility did not change in response to 1 min of morphine (10 µM) treatment, but after 10 min of morphine treatment, tracks shifted towards immobility. Closer inspection of mobile (*α* > 0.27) tracks revealed that differences in MSD exist between experimental conditions even within the mobile subpopulation of receptors, and that this difference in MSD can be explained by an increase in confined dwell times of receptors treated with DAMGO for ten minutes. In contrast, MORs treated with DAMGO for one minute show a decrease in confined dwell times, and blockade of G-protein binding with pertussis toxin abrogates this decrease in confined period dwell times. Finally, CCP colocalization occurred similarly across all mobility states inspected so that a single mobility state could not fully account for colocalization of FLAG-MORs with CCPs.

Some studies have shown that Qdots can reduce molecule mobility and alter molecular interactions^28,36^. In this study, great care was taken during the labeling approach to avoid receptor crosslinking and to ensure that the MOR was still undergoing G-protein coupling and recruitment to GFP-CLC when expected. Further, experiments with Dermorphin-488 and correlation of MSD and fluorescence intensity ensured that crosslinking likely plays a minimal role in these experiments. However, it is still possible that Qdots have slowed diffusion compared to an unlabeled MOR. Nevertheless, we were interested in the dynamic changes in response to drug application as opposed to basal receptor mobility and changes in mobility were observed in response to drug. Thus, our observations still stand even if Qdots have slowed mobility.

Previous studies of MOR mobility in response to agonist treatment have examined mobility at a minimum of ten minutes post-drug, although acute signaling processes occur much faster than this. In AtT20 cells, the receptor has already undergone G-protein coupling, desensitization, and internalization at this time point^13^. Thus, we were interested in understanding how MOR mobility is changed before internalization has begun and while the process of signaling and desensitization are occurring. The marked difference in receptor mobility depending upon duration of agonist exposure reveals that mobility is likely dependent on receptor interactions with intracellular partners that play distinct roles depending upon the time of drug application. The difference in alpha value distribution between morphine (1 min) and DAMGO (1 min) is particularly interesting, as these agonists have distinct signaling characteristics. The fact that DAMGO, but not morphine, increased the mobile fraction of receptors at 1 min may be attributable to the relatively lower efficacy of morphine compared to DAMGO for recruiting G-proteins at the MOR^37^. However, it is also possible that morphine’s lower ability to recruit other effectors such as specific kinases or β-arrestins could play a role in the lack of shift towards mobility at 1 min.

Upon inspection of mobile trajectories in free and confined states, it was initially surprising to note that acute DAMGO treatment resulted in a decrease in the time that mobile tracks spent confined. However, a recent study by Sungkaworn et al^38^ revealed that G-protein coupled receptors and their respective G-proteins couple at specific hot spots at least partially defined by actin networks^38^. It is possible that signaling receptors in AtT20 cells quickly hop between G-protein hot spots after activation, thus decreasing their confined dwell times. This possibility is supported by the fact that PTX increased confinement periods compared to baseline levels, potentially suggesting that ligand-bound receptors are recruited to hot spots but cannot leave unless they have encountered a G-protein. However, it should be noted that the change in dwell times between 1 min DAMGO and no drug conditions is small, therefore binding to other signaling partners likely also plays a role in how long a receptor remains confined or free.

Despite the difference in acute mobility shifts, both DAMGO and morphine increased the immobile fraction of receptors after 10 min of exposure. Morphine is known to be poor at inducing receptor internalization compared to DAMGO^39,40^, yet track mobility still decreased after 10 min of exposure. At this time point, both DAMGO and morphine can cause receptor desensitization. Thus, the shift towards receptor immobility likely reflects interactions related to desensitization. Kinases such as protein kinase C are known to be involved in receptor desensitization and have been found to restrict the distribution of the MOR in cell membranes^3^. It is possible that other kinases, association with lipid rafts^41^, or binding to structural proteins ^42^ could contribute to immobility as well. Future studies using receptors that do not desensitize, such as non-phosphorylatable mutants^16,43^, could best parse apart the contribution of desensitization and G-protein coupling to transient confinement in mobile MOR trajectories.

The present study shows increased colocalization of MORs with clathrin after 10 min of DAMGO exposure, consistent with the ability of DAMGO to strongly induce internalization. Morphine-induced colocalization was not examined because of its low ability to induce internalization and because we were unable to attribute any particular mobility state to receptor capture into CCPs. Our finding that colocalization with CCPs cannot fully account for any single mobility state suggests that this may be true at other effectors as well. Therefore, the immobile fraction of receptors observed after 10 min DAMGO treatment is likely due to the contribution of several effectors.

Because CCP colocalization was not associated with a particular mobility population that we identified, it is unlikely that single-particle tracking of the MOR alone would be a good proxy for distinct receptor signaling states. However, concurrent tracking of the MOR with its signaling effectors may be useful as a tool for studies of receptor/effector interaction. For example, concurrent single-particle tracking of the MOR, G-proteins, and β-arrestin or CCPs would allow for screening of ligand bias within the same cell with high temporal resolution and individual receptor sensitivity.

## Materials and Methods

### Cell Culture and Transfection

AtT20 cells stably expressing the FLAG-MOR (provided by Dr. MacDonald Christie, University of Sydney) were maintained at 37°C/5% CO_2_ in Dulbecco’s Modified Eagle Medium (DMEM) supplemented with 10% fetal bovine serum (ATCC), L-Glutamine (2 mM L-alanyl-L-glutamine), and 1% penicillin/streptomycin. Once cells reached confluence they were exposed to 0.25% trypsin-EDTA (Gibco) and re-plated at lower density. Cells were maintained for no more than 12 passages beyond their original plating.

N-terminally tagged GFP-clathrin light chain (GFP-CLC) was transfected into cells using 1:500 Lipofectamine (Invitrogen), 1:100 Opti-MEM (Gibco), and 1:1000 plasmid into 2 mL of culture media. Cells were incubated overnight, and all cells were imaged within 24 hours of transfection in order to avoid problems with overexpression. For experiments in which G-protein activity was inhibited, cells were treated with 100 ng pertussis toxin (Thermo Fisher) overnight. Cells were imaged within 24 hours of toxin treatment.

### Live-Cell Labeling of the MOR

To prepare for imaging, AtT20 cells were diluted and plated on glass-bottom dishes (MatTek, Ashland, MA) containing the same supplemented DMEM solution as described above. Cells were imaged on the fourth day after plating. Labeling was performed immediately before imaging. Cells were rinsed multiple times with a saline solution containing: 35 mM KCl, 120 mM NaCl, 1 mM CaCl_2_, 25 mM HEPES, 10 mM glucose, and pH was adjusted to 7.4 (NaOH). After thoroughly washing to remove DMEM, the plates were filled with saline containing 1% (w/v) bovine serum albumin (BSA, Sigma A7030) and incubated for 10 min at 37°C. The cells were incubated for 5 min at 37°C in the presence of a biotinylated anti-FLAG antibody (final concentration 1 µg/mL, BioM2 Anti-FLAG, Sigma F9291). After multiple rinses with saline (still containing 1% BSA) to remove unbound antibody, the cells were then incubated for 8 min at 37°C in the presence of streptavidin-coated quantum dots (1:10000, final concentration of 100 pM, Streptavidin Qdot 565, Life Technologies Q10133MP; or for two-color TIRF imaging Streptavidin Qdot 655, Life Technologies Q10121MP). The cells were rinsed multiple times with saline lacking BSA to remove unbound SA Qdot and BSA from the culture dish. In the 10 min DAMGO condition and Naloxone condition, 10 µM DAMGO or 1 µM Naloxone, respectively, was applied immediately after labeling. Drug was applied 10 min after the final wash on the microscope stage in the 1 min (10 µM) DAMGO condition. In experiments where morphine was used as the agonist rather than DAMGO, morphine (10 µM) was applied as described above for DAMGO. Many cells did not exhibit Qdot fluorescence upon imaging, so only those that had a moderate amount of labeling were chosen for imaging.

### Confocal Microscopy

Single-channel imaging was carried on a spinning disk confocal microscope (Olympus IX83, Olympus UPlanSApo 100x/1.40 oil objective, Yokogawa CSU-X spinning disk, Andor iXon Ultra 897 EMCCD camera) equipped with a temperature control unit (INU Stage Top Incubator, Tokai Hit, Shizuoka-ken, Japan). All culture dishes were kept in the chamber at 37°C for 10 min before images were acquired. This was done to ensure that all dishes were imaged at the same time after labeling. Qdot 565 was excited using a 488 nm laser and acquired with an emission filter (600/50). Videos were acquired at a rate of 20 frames/s.

### TIRF Microscopy

GFP-CLC was found to bleach rapidly using the spinning disk confocal, so for concurrent imaging of GFP-CLC and the MOR, total internal reflection fluorescence (TIRF) microscopy was performed on a Nikon Eclipse Ti fluorescence microscope equipped with a Perfect-Focus system, AOTF-controlled 488 and 647 nm diode lasers, a 512×512 Andor iXon EMCCD DU-897 camera, and Plan Apo TIRF 100, NA 1.49 objective. Temperature was maintained at 37°C using Zeiss stage and objective heaters. Qdot 655 was used instead of Qdot 565 to avoid bleed-through into the 488 channel for imaging CLC-GFP. To avoid analyzing Qdots that occasionally were stuck to the culture dish glass, trajectories with MSDs characteristic of glass-stuck particles were removed before analysis. A sample of glass-stuck Qdots were imaged and found to have mostly MSDs below 0.0165 μm^2^ (Fig. S3A). Therefore, all trajectories with an MSD less than 0.0165 μm^2^ (measured up to a lag time of 2.5 s) were excluded from the data prior to analysis.

### Image Processing and Analysis

Images were background-subtracted in ImageJ software. A Gaussian kernel filter was then applied to the images using a standard deviation of 0.8 pixels. After processing, Qdot-labeled MORs were detected and tracked using the u-track algorithm in MATLAB as previously described^44^. Detection of CLC-GFP puncta was also performed using the u-track algorithm. MOR trajectories less than 60 frames in length were excluded from further analysis.

Trajectories were analyzed in terms of the time-averaged mean square displacement (TA-MSD) using algorithms written in MATLAB. For an individual trajectory the TA-MSD is obtained by averaging over the time series,

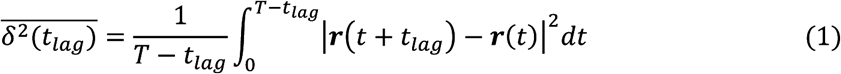

where 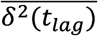 is the TA-MSD, ***r***(*t*).is the two-dimensional position of the particle at time *t, t*_*lag*_ is the lag time (the time over which the displacement is computed), and *T* is the duration of the trajectory. For normal diffusion processes, the MSD scales linearly in lag time, namely in two dimensions 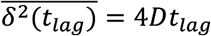, where *D* is the diffusion coefficient. However, measurements in live cells often exhibit anomalous diffusion, which manifests as a deviation from this simple law^20-22^, and is characterized by a non-linear scaling of the MSD, 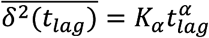 where α is the anomalous exponent and *K*_*α*_ is the generalized diffusion coefficient which has units of cm^2^/s^α^. Processes with 0 < α< 1 are considered subdiffusive, and those with α > 1 are considered superdiffusive. Detection uncertainty increases the MSD by a constant value. Given a standard deviation σ of the detected position in both *x* and *y* direction due to uncertainty in the localization, the MSD is then

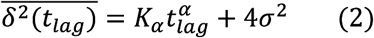

In order to obtain α and *K_α_* from individual trajectories, we first obtained an average σ of 0.02 with the u-track algorithm and subtracted 4σ^2^ to obtain a static error-corrected MSD (see Fig. S3B for a histogram of localization uncertainties, σ). Then we perform a linear regression in log-log plot to find α and *K_α_*.

To separate mobile tracks into free and confined portions, recurrence analysis was performed as described previously^30^. Briefly, a circle is constructed equal to the diameter of two consecutive points in a trajectory, and the number of subsequent visits by the particle into this circle was calculated. Portions of tracks with subsequent visits ≥ 11 were classified as confined portions of mobile trajectories, and those that did not reach this threshold were classified as free portions. This threshold was determined by identifying the minimum number of visits at which most immobile (α < 0.27) trajectories were classified as confined.

To calculate the colocalization and distance of the MOR to CCPs, spatial coordinates of GFP-CLC puncta were determined using the detection output in u-track. The coordinates of GFP-CLC puncta were then compared to the coordinates of MOR tracks files at each frame. MOR coordinates that were located within 3 pixels of the closest GFP-CLC pit center (0.48 µm) were considered to be colocalized.

### Statistical Analysis

Data sets were compared using an unpaired Kruskall-Wallis one way ANOVA with post hoc tests performed with Bonferroni correction, and *p* values of less than 0.05 were considered significant. Prism was used to perform statistical tests as well as to obtain descriptive statistics. Compiled data are shown as the mean ± SEM, or as histograms.

## Supporting information

## Acknowledgments

The authors thank Dr. Michael Tamkun and his group for providing expertise and equipment necessary to carry out these studies. In particular, Emily Maverick and Dr. Laura Solé provided critical assistance in imaging and single-particle tracking. Additionally, we thank Dr. Sanaz Sadegh, Kanti Nepal, and Xinran Xu for providing technical assistance with MATLAB codes used for data analysis and Drs. John T Williams and Seksiri Arttamangkul for providing labeled compounds. Funding for these studies was provided by National Institutes of Health grants R01DA032562 (S.T.H.) and F31DA035586 (R.L.P.), as well as National Science Foundation grants DGE-1321845 (M.J.M) and 1401432 (D.K.).

## Author Contributions

M.J.M, R.L.P, D.K., and S.T.H designed experiments. M.J.M. and R.L.P performed experiments. D.K. provided MATLAB codes used in analysis, and M.J.M, R.L.P., and D.K. analyzed data. M.J.M., R.L.P, D.K., and S.T.H. wrote the manuscript.

## Additional Information

The authors have no conflicts of interest related to this work.

