## Supplementary material for "Temporal dependence of shifts in mu opioid receptor mobility at the cell surface after agonist binding observed by single-particle tracking"

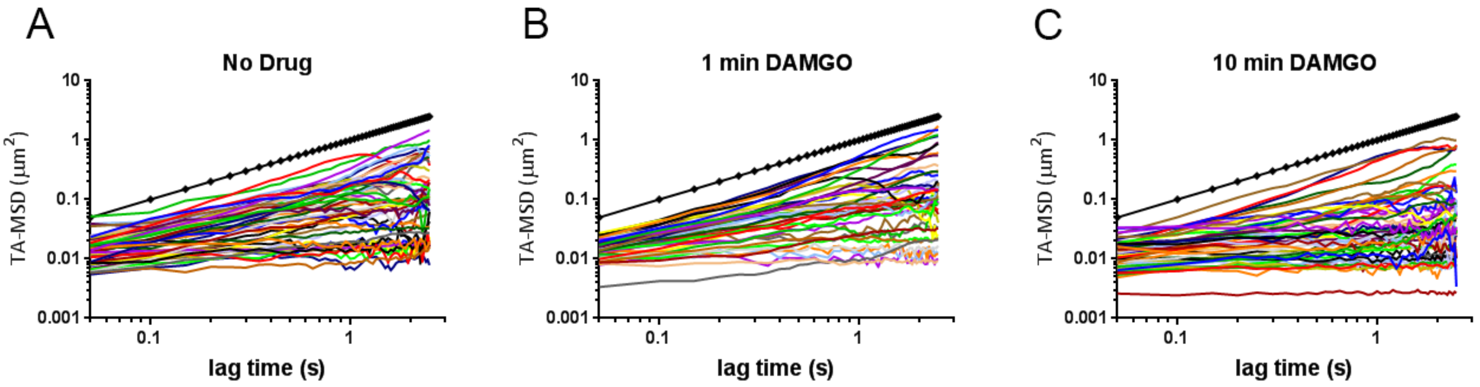


**Figure S1. Individual MSDs of randomly chosen trajectories.** Log-log plots of time averaged MSDs from individual tracks in the **A)** no drug, **B)** 1 min DAMGO, and **C)** 10 min DAMGO experimental conditions. Most trajectories are subdiffusive, as they lie below the slope of a simulated track of $\alpha$ = 1 (black dotted line).

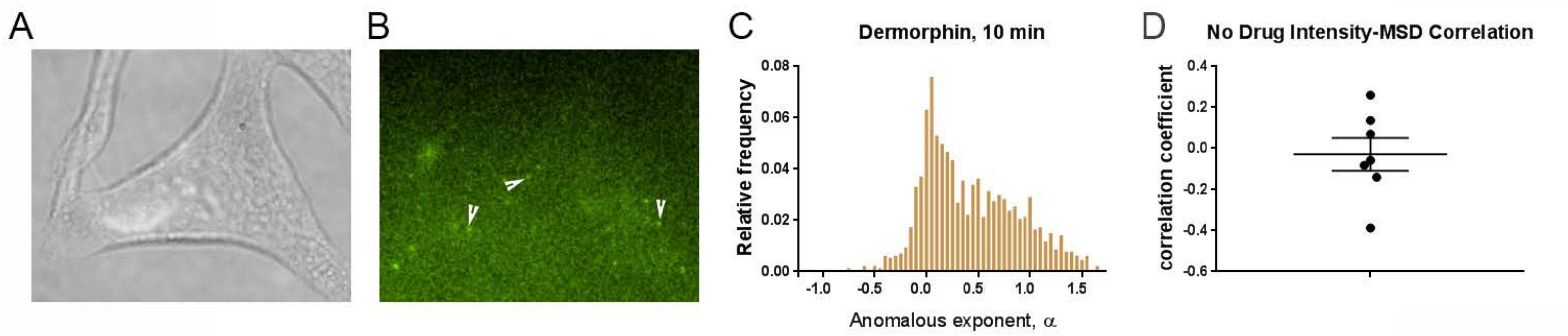


**Figure S2. Single particle tracking after 10 min of Dermorphin-488 (60 pM) application reveals that the immobile population of MORs is not due to antibody-mediated crosslinking or Qdot hindering of mobility. A.** DIC image of an AtT20 cell labeled with Dermorphin-488. **B)** The same cell is shown under fluorescence, and arrows indicate MORs labeled with a single Dermorphin-488 conjugate. **C)** Distribution of $\alpha$ values after tracking of MOR-Dermorphin-488 conjugates (n = 5 cells, 1266 tracks) incubated for 10 min in the presence of 60 pM Dermorphin-488. The distribution of $\alpha$ values is similar to those observed with Qdot tracking, and the fraction of $\alpha < 0.27$ is 0.44 ± 0.08, similar to MOR-Qdots in the 10 min DAMGO condition (0.45 ± 0.12). Values below 0 are likely due to errors made during tracking caused by the low signal to noise ratio with this labeling approach. **D)** Distribution of MSD vs. fluorescence intensity correlation coefficients for individual cells in the no drug condition, with a mean correlation coefficient of -0.03 ± 0.08.

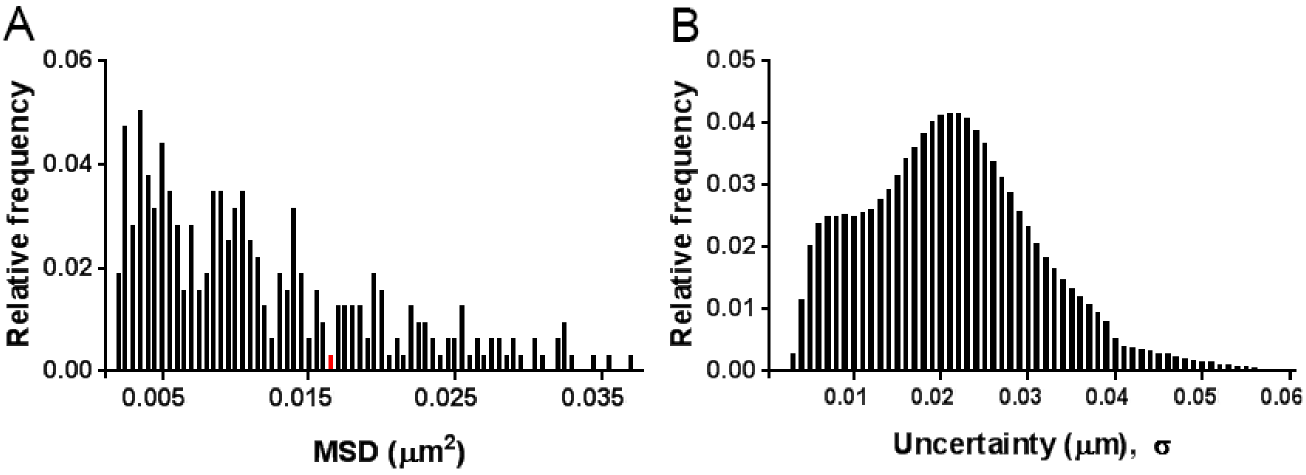


**Figure S3. Data for trajectory corrections. A)** Frequency histogram of MSD for glass-stuck Qdot-655 tracked without cells on the coverslip. The cutoff of 0.0165 $\mu m^2$ is highlighted in red. Most MSDs are less than this cutoff. **B)** Histogram of localization uncertainties, $\sigma$, for each detected particle of a representative subset in the no drug condition. An average $\sigma$ of 0.02 $\mu m$ was used for uncertainty correction of tracks.
